# 15-Hydroxyeicosatetraenoic Acid and GPR39 Together Orchestrate Coronary Autoregulation: A Comprehensive Metabolomic Analysis

**DOI:** 10.64898/2025.12.23.696315

**Authors:** D. Elizabeth Le, Masaki Kajimoto, Yan Zhao, Carmen Methner, Zhiping Cao, Jessica Minner, Agostino Cianciulli, Teresa Semeraro, Iuni Margaret Laura Trist, Jessica Franchi, Chiara Marcheselli, Alberto Parazzoli, Fabrizio Micheli, Sanjiv Kaul

## Abstract

**Background:** Coronary autoregulation is the ability of the normal heart to maintain constant coronary blood flow (CBF) over a wide range of coronary driving pressures (CDP). Despite being vital for survival, the mechanism of coronary autoregulation is unknown. We hypothesized that GPR39, present in vascular smooth muscle cells, together with its endogenous agonist 15- hydroxyeicosatetraenoic acid (15-HETE) orchestrate coronary autoregulation.

**Methods:** We created coronary stenoses of varying degrees in open-chest, anesthetized dogs where we measured CBF and CDP. In a subset of animals, coronary venous blood was sampled for eicosanoid, adenosine, endothelin-1, polyunsaturated fatty acids, and prostaglandins levels. Stenoses were recreated during intravenous administration of VC108, a specific GPR39 antagonist and systemic, pulmonary, and coronary hemodynamics measured.

**Results:** GPR39 was identified in coronary arterioles by immunohistochemistry and in heart tissue by western blot. In-vivo, 15-HETE correlated the best (r^2^=0.4851, p=0.0144) with CDP over the autoregulatory range using a linear mixed-effects model. Prior to administration of VC108, CBF did not change within the autoregulatory range. VC108 had no effect on systemic and pulmonary hemodynamics but increased CBF (p=0.02 versus vehicle) by decreasing coronary microvascular resistance (p=0.01 versus vehicle), indicating that GPR39 participates in control of normal coronary vascular tone. With VC108, coronary autoregulation was abolished and CBF became CDP dependent (r^2^=0.9562, p=0.0039).

**Conclusion:** GPR39 and its endogenous agonist 15-HETE together orchestrate coronary autoregulation when CDP is reduced. These novel findings provide a mechanism for coronary autoregulation and could direct pharmacological treatment of various coronary syndromes in humans.

**Clinical Perspectives:** 

**What is new?:** The new finding from this study is that the vasoconstrictor, 15- hydroxyeicosatetraenoic acid (15-HETE), acting via the G-protein coupled receptor 39 (GPR39) participates in coronary arteriolar tone, and when perfusion pressure falls because of coronary stenosis, 15-HETE levels decrease causing vasodilation to maintain resting coronary blood flow.

**What are the Clinical implications:** This mechanism, termed coronary autoregulation, causes patients with coronary artery disease to remain asymptomatic at rest until coronary stenosis is very severe.

Pharmacological inhibition of GPR39 causes selective coronary vasodilation and could be used to treat coronary artery disease.

## Main

Coronary autoregulation refers to the ability of the normal heart to maintain constant coronary blood flow (CBF) over a wide range of coronary driving pressures (CDPs) provided cardiac work remains constant. Lowering CDP results in vasodilation of the resistance arterioles present within the heart while raising CDP cause vasoconstriction of these vessels, thus maintaining constant CBF.^1–4^ Autoregulation is a teleologically protective mechanism that is meant to keep blood flow constant to vital organs despite wide alterations in systemic pressures and to ensure a constant capillary hydrostatic pressure necessary for cell survival.^5^ In large animals and humans, the CDP range where CBF remains constant is between approximately 45 to 120 mm Hg.^1–4^

Despite the relevance of coronary autoregulation to survival, its mechanism is unknown.^6^ The eicosanoid 20-hydroxyeicosatetraenoic acid (20-HETE) has been implicated in cerebral autoregulation.^7^ Eicosanoids form a wide variety of lipid-based signaling molecules present in various forms and concentrations in different tissues.^8^ Although some eicosanoids are vasoactive, their role in coronary autoregulation has not been examined. Further, while single mediators like adenosine have been investigated as a putative mechanism of coronary autoregulation,^9,10^ a comprehensive metabolomic analysis of all possible vasoactive compounds that could influence resting coronary vascular tone and coronary autoregulation has not been performed.

We reported the presence of a G-protein coupled receptor (GPR39) in the vascular smooth muscle cells (VSMCs) of coronary arterioles in the mouse heart.^11^ We showed that two eicosanoids are endogenous ligands for GPR39: 15-hydroxyeicosatetraenoic acid (15-HETE) is the agonist, and 14,15-epoxyeicosatrienoic acid (14,15-EET) is the antagonist. GPR39 stimulation by 15- HETE activates phospholipase C through the Gαq pathway that cleaves membrane bound inositol phosphatidylinositol 4,5-bisphosphate (PIP2) into inositol 1,4,5-trisphosphate (IP3) and diacylglycerol (DAG). DAG remains in the cell membrane while IP3 binds to the IP3 receptors on the endoplasmic reticulum, releasing Ca^++^ into the cytosol, causing cell contraction.^11^ Ca^++^ also acts as a second messenger causing extracellular signal-regulated kinase (ERK) phosphorylation leading to additional downstream effects.^11^

Both 15-HETE (vasoconstrictor) and 14,15-EET (vasodilator) act through GPR39 to maintain coronary vascular tone.^11^ Consequently, we hypothesized that the levels of 15-HETE and 14,15-EET, acting on GPR39, will determine coronary vascular tone depending on the CDP. We postulated that the receptor and its ligands together orchestrate coronary autoregulation.

## Methods

A canine model was selected for the autoregulation study because coronary physiology can be reliably studied in these animals and because most of the classical papers on coronary autoregulation were performed in this model. The data that support the findings of this study are available from the corresponding author upon reasonable request. Results are reported according to ARRIVE guidelines. The study protocols were approved by the Institutional Animal Care and Use Committee at the Oregon Health & Science University and conformed to the National Institutes of Health guidelines for animal research.

### Animal Preparations

For the autoregulation study, 35 adult male mongrel dogs (30-35 kg) were studied. Females were excluded because of possible pregnancy and uncertainty of timing in the ovulation cycle. Dogs were sedated with intramuscular injection of hydromorphone (0.1-0. 2 mgꞏKg^-1^) and then induced with intravenous administration of ketamine (2-5 mgꞏKg^-1^) and midazolam (0.1-0.4 mgꞏKg^-1^). They were then intubated and ventilated (Integra SL with AV-S ventilator, Avante Health Solutions, Louisville, KY, USA) with isoflurane gas (0.5-1.0%) and medical-grade room air. Constant-rate infusion anesthesia was maintained with intravenous administration of ketamine (0.3-0.5 mgꞏKg^-1^ bolus followed by an infusion of 0.3-1.5 mgꞏKg^-1^hr^-1^), fentanyl (0.001-0.003 mg bolus followed by an infusion of 0.05-0.07 µgꞏKg^-1^min^-1^), and a very low dose lidocaine infusion (0.6-6.0 mgꞏKg^-1^hr^-1^ IV). These infusion rates of the drugs were adjusted throughout the experiment to optimize anesthesia without affecting coronary reactivity. The mean coronary hyperemic response after a 20 s occlusion was 3-fold, which is normal in open-chest dogs indicating that anesthesia did not affect vasomotor tone.

Heart rate, blood pressure, O_2_ saturation, end-tidal CO_2_, and temperature were continuously monitored (Advisor® Vital Signs Monitor, Surgivet, Norwell, MA, USA). A 6F sheath was inserted in the right femoral artery for arterial pressure measurement. 7F catheters were inserted into the right femoral, right atrium (RA), and peripheral veins for administration of drugs and fluids and measurement of pressure. In a subset of animals, a 7F sheath was inserted for placement of a balloon-tipped catheter (Edwards Lifesciences, Santa Ana, CA, USA) for measurement of pulmonary artery (PA) pressures. In the same dogs a micromanometer-tipped catheter (Millar Instruments, Inc., Houston, TX, USA) was inserted into the left ventricular cavity to measure left ventricular ΔP/Δt.

A left lateral thoracotomy was performed, and the heart was suspended in a pericardial cradle. The left anterior descending (LAD) coronary artery was dissected free from surrounding structures in its proximal and mid portions. An ultrasonic time-of-flight flow probe (Series SC, Transonics, Ithaca, NY, USA) was placed on the proximal segment of the artery and connected to a digital flow meter (Model T206, Transonics). A screw occluder was positioned around the mid-portion of the artery to create coronary stenoses of varying degrees. A 20-gauge polyethylene catheter was inserted into the distal portion of the artery to measure pressure. A similar catheter was inserted in the anterior cardiac vein for obtaining blood samples. CDP was calculated as the difference between the mean distal LAD and RA pressures. Coronary microvascular resistance (MVR) was derived by the mean CDP/CBF ratio.

### Hemodynamic Measurements

All catheters and flow meters were interfaced with a multi-channel recorder (PowerLab, AD Instruments, Inc., Colorado Springs, CO, USA). Phasic and mean CBF and systemic pressures were acquired digitally and displayed on-line on an iMac (Apple Inc., Cupertino, CA, USA). All data were analyzed off-line using LabChart 6.

### Mass Spectroscopy

The details for mass spectroscopy measurements of lipids and adenosine have been published before.^12,13^ For measurement of eicosanoids and prostaglandins, blood was drawn in a stop solution containing 8 mg butylated hydroxytoluene and 80 mg triphenylphosphonium mixed with 80 mg indomethacin in 40 mL methanol solution. The samples were centrifuged, placed on ice and sent for analysis.

Measurements were made using MS/MS with a 5500 Q-TRAP hybrid/triple quadrupole linear ion trap mass spectrometer (Applied Biosystems, Waltham, MA, USA) with electrospray ionization (ESI) in negative mode. The mass spectrometer (MS/MS) was interfaced to a Shimadzu (Columbia, MD, USA) SIL-20AC XR auto-sampler followed by 2 LC-20AD XR LC pumps and analysis on an Applied Biosystems/SCIEX Q5500 instrument. The instrument was operated with the following settings: source voltage -4000 kV, GS1 70, GS2 70, CUR 45, TEM 750 and CAD gas MED. The gradient mobile phase was delivered at a flow rate of 0.5 mL/min, and consisted of two solvents, A: 0.05% acetic acid in water and B: acetonitrile. The LC column used was a Kinetex Phenyl-Hexyl 50×2.1 mm 2.6 µM. The initial concentration of solvent B used was 20%, this was held for 0.5 minutes, increased to 60% B over 10.5 minutes, increased to 95% B over 1 min, held at 95% B for 1 min, and then returned to initial conditions over 0.1 min, followed by re-equilibration for 1.9 min for a total run time of 15 min. The column oven was set to 50°C, autosampler to 15°C with needle rinse before and after each injection. Data was acquired using Analyst 1.7.1 and analyzed with Multiquant 3.03.

For arachidonic acid and other polyunsaturated fatty acids, analysis was performed using an Agilent 7890B GC (Santa Clara, CA, USA) with a 5977A MS detector with an autosampler and split/splitless injector operated in split mode. The column was an Agilent DB-FastFAME column (30 m, 0.25 mm id, 0.25 µm film thickness). Helium was the carrier gas at a flow rate of 1 mL/min. The injection port and auxiliary heater were maintained at 245°C. A 1 µL sample was injected in split mode (1:10) at an initial oven temperature of 150°C, held for 0.5 min and then increased at 15°Cꞏmin^-1^ to 200°C, followed by 25°Cꞏmin^-1^ to 240°C, held for 2 min and then returned to 50°C. The MS was operated at a source temperature of 230°C and a MS quad temperature of 150°C in positive electron impact mode. The GC-MS data were acquired using enhanced MassHunter software. Calibration curves for quantification were generated from peak area ratios for authentic standard:internal standard for calibrator samples.

For adenosine, blood was obtained in stop solution containing dipyridamole, sodium EDTA, erythro-hydroxy-nonyladenine, α,β-methylene-adenosine 5’-diphosphatenucleotidase, and heparin.^14^ The sample was centrifuged, placed on ice, and sent for analysis. Plasma was analyzed using MS/MS with a 5500 Q-TRAP PLUS hybrid/triple quadrupole linear ion trap mass spectrometer (Applied Biosystems) with electrospray ionization (ESI) in positive mode. The tandem MS/MS was interfaced to a Shimadzu SIL-20AC XR auto-sampler followed by 2 LC- 20AD XR LC pumps. The MS instrument was operated with the following settings: source voltage 5500 kV, GS1 60, GS2 560, CUR 35, TEM 750, and CAD gas MED. Compounds were infused individually and MS/MS instrument parameters optimized for each MRM transition. Those used for quantification were *m/z*=268→136 for adenosine with a dwell time of 100 msec, DP= 75, EP=10, CXP=15 and CE=25 and *m/z*=273.2→136 with a dwell time of 100 msec, DP= 75, EP=10, CXP=15 and CE=25. Data was acquired using Analyst 1.7.3 and analyzed with Multiquant 3.03. For VC108 measurements, a 12-point calibration curve and quality control samples were prepared in blood diluted 1:4.8 in 0.1N Hepes buffer in micronic tubes., 20 µL of which was transferred into micronic tubes containing 80µL 0.1N Hepes buffer. Calibration samples, quality control samples, and analysis samples were extracted by adding 400µL CH3CN containing IS (Rolipram) 20 ng/mL.

After vortex and centrifugation (3000 rpm/10 min), samples supernatants were transferred to a 96 well plate and diluted (450µL CH3CN/aq=37/63 + 50µL supernatant). Samples were injected using an Agilent 1290 Infinity II UPLC system (Santa Clara, CA, USA) and separated on a Acquity UPLC BEH C18 (30×2.1mm, 1.7µm, Milford, MA, USA) connected directly to the TurboIonSpray source of an AB Sciex 6500QTrap mass spectrometer (Framingham, MA, USA). Mobile Phase A was 0.1% (v/v) formic acid in water and Mobile Phase B was 0.1% (v/v) formic acid in acetonitrile. The flow rate was 1 mL/min and a generic gradient of 5% B to 95% B in 1.3min was applied.

### ELISA for Endothelin-1

Plasma samples were analyzed for endothelin-1 using an ELISA kit (Catalog# ADI-900- 020A, Enzo, Farmingdale, NY, USA) as described in the manufacturer’s manual. The absorbance was measured at 450nm using a 96-well plate reader (Molecular Devices, FlexStation 3, San Jose, CA, USA).

### Immunohistochemistry

After euthanasia, hearts were removed from 3 dogs that had received vehicle during the experiments. Tissues were fixed with fresh 4% paraformaldehyde and cut by cryostat into 20 µm sections that were mounted on superfrost glass slides, washed 3 times with PBS containing 4% goat serum, 1% bovine serum albumin, and 0.3% Triton-100 for 90 at room temperature. Sections were then incubated overnight with rabbit anti-GPR39 primary antibody (1:200, Cat #AP19112b, Abgen, San Diego, CA, USA). After washing 3 times with PBS, sections were intubated with the secondary antibody (1:200, Alexa 488-conjugated donkey anti-rabbit (Cat #A21206, Thermo Fisher Scientific, Waltham, MA, USA) for 2 h at room temperature. Sections were then washed 3 times in PBS and placed in autofluorescence reduction solution (10 mM CuS04, 50 nM ammonium acetate) for 30 min followed by 2 min incubation with trihydrochloride trihydrate (1:4000 Cat #33342, Invitrogen, San Diego, CA, USA) to stain nuclei. The sections were mounted using Fluoromount G® (Cat #0100-01, Sothern Biotech, Birmingham, AL, USA). Digital images of the sections were acquired at 20x with Nikon Imaging System (Nikon Eclipse Tie-A1RSi, Tokyo, Japan).

### Western Blot

Heart tissues from 3 dogs were homogenized in liquid nitrogen and lysed in RIPA buffer (Thermo Fisher Scientific, Cat. # PI89900) with protease and phosphatase inhibitor (Pierce, Thermo Fisher Scientific, Cat. # A32959). The homogenates were then centrifuged at 14,000 rpm for 20 min at 4°C. The Pierce BCA Protein assay kit (Thermo Fisher Scientific, Cat. # 23225) was used to measure the concentration of protein lysate in samples. Whole heart protein (20 µg per lane) was separated on 4%–12% SDS-polyacrylamide gels and transferred to polyvinylidene difluoride membranes. Total protein staining was performed using Licor Bio Revert® 700 total Protein Stain (Licor Bio, Cat. #926-11016). The membranes were blocked with Licor Intercept® TBS blocking buffer (Licor Bio, Cat. #927-60003) for 60 min at room temperature followed by primary antibodies blocking buffer with 0.02% Tween-20: 1:500 rabbit anti-GPR39 antibody (Novus Biologicals, #NLS139). After 1 h incubation with secondary antibody (IRDye® 800CW Goat anti-Rabbit IgG Secondary Antibody, 1:5000, Licor Bio, Cat. # 926-32211) at room temperature, the membranes were imaged using Licor Odyssey Imager with Image Studio software.

### Study Protocols and their Rationale

We used 3 groups of dogs (Figure 1). Group 1 comprised 7 dogs in which coronary stenoses of different degrees were created within the autoregulatory range and eicosanoid levels from the anterior cardiac vein were measured at each stage. Our aim was to determine whether eicosanoids levels changed during decreases in CDP, which would allow us to further explore our hypothesis. Based on the results from Group 1 dogs we proceeded with experiments in the 28 Group 2 dogs (Figure 1). By then we had access to a specific GPR39 inhibitor, VC108,^15^ that had been found to increase CBF in rodents.^16^ To test off-target VC108 binding, a 131 panel of assays including receptors, enzymes, channels. and transporters was performed at Eurofins Scientific (Luxembourg, Supplemental methods).

**Figure 1:**
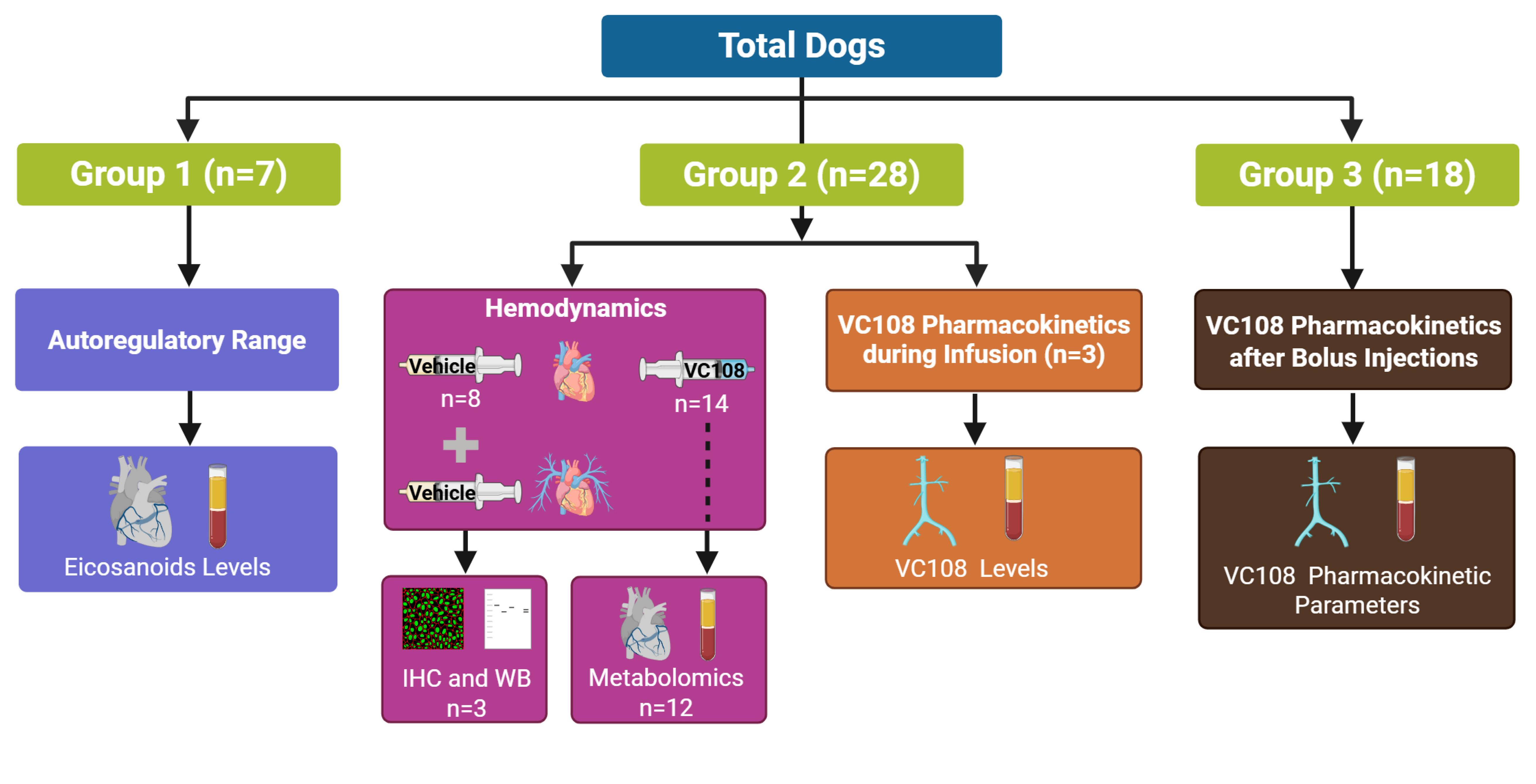
Breakdown of the animals used in the study and protocols they underwent. IHC=immunohistochemistry; WB=western blot. Created in https://Biorender.com.

We created stenoses of varying severities (including flow limiting) in 22 of these dogs to demonstrate the presence of coronary autoregulation, and in 12 of them we collected anterior cardiac venous blood at each stage for metabolomic analysis: adenosine, endothelin-1, polyunsaturated fatty acids (including arachidonic acid and prostaglandins). We then administered VC108 (n=14) or vehicle (n=8) in these animals and added 3 additional dogs to the vehicle group to make the total =11, and measured systemic, pulmonary and coronary hemodynamics 30 min later. Vehicle comprising 10% DMSO was administered as a 5-8 mL IV bolus over 30 s followed by a continuous infusion (120 mL·hr^-1^). Three of these dogs receiving vehicle were used for immunohistochemistry and western blot analysis for GPR39. VC108 suspended in 10% DMSO was administered as a 1.5 mg IV bolus over 30 s followed by a continuous infusion (0.6 mgꞏKg^-1^ꞏhr^-1^). This dose was calculated based on >90% receptor occupancy. We then recreated stenoses of varying severities in the 14 dogs receiving VC108 to demonstrate abolishment of coronary autoregulation. Finally, we used 3 additional dogs in which blood levels of VC108 were measured every 10 min for an hour.

An additional group of Beagle dogs (Group 3) were studied at Aptuit for VC108 pharmacokinetics following daily intravenous administration for 14 days. VC108 was given to 18 dogs (3/sex/group) at 2, 6 and 20 mgꞏkg^-1^ once daily for 14 days by intravenous (bolus) administration. Dose volume administered per day was 2.5 mLꞏkg^-1^. Pharmacokinetic parameters (drug exposure) were evaluated on Days 1 and 14 (Supplemental Methods).

At the end of the experiment, with animals still under deep anesthesia, 1 mLꞏ4.55 kg^-1^ of Euthasol (sodium pentobarbital 390 mgꞏmL^-1^ and sodium phenytoin 50 mgꞏmL^-1^) was administered intravenously in accordance with the June 2020 American Veterinary Medical Association guidelines on euthanasia that is approved by the Institutional Animal Care and Use Committee at our institution.

### Statistical Methods

Data were expressed as mean±1 standard error of the mean (SEM). Because observations were repeated within animals (multiple stenosis stages per dog), eicosanoid versus CDP levels in the Group 1 dogs were analyzed using linear mixed-effects models (LMMs) with CDP as a continuous predictor and a random intercept for dog; fixed-effect p values used the Satterthwaite approximation and were Bonferroni-corrected across the eicosanoid panel. Then a multivariable LMM was fit with only significant predictors and a likelihood ratio test was used to compare the model with multiple predictors and the univariable model containing just 15-HETE. Comparison of hemodynamic data between vehicle and VC108 in Group 2 dogs was performed using unpaired t test. Systemic and coronary hemodynamics across stenosis stages, prior to and during VC108 administration in the Group 2 dogs, were analyzed using LMMs with stenosis stage as a fixed factor and a random intercept for dog, with Tukey-adjusted pairwise comparisons between stages. CDP versus all other cardiac venous metabolites in the 12 Group 2 dogs sampled prior to VC108 administration was analyzed using LMMs with continuous CDP and a random intercept for dog, with false-discovery-rate correction across the metabolite panel. Data before and during VC108 administration were compared using a lineLMM with a condition-by-stage interaction and a random intercept for dog. Aggregate CDP versus CBF data in the 22 dogs prior to VC108 administration were fit to a one phase decay function. Linear regression for performed for the aggregate CDP versus CBF data in 14 Group 2 dogs during VC108 administration. Significance was determined at P<0.05 (two-sided).

## Results

Figure 2 illustrates examples of IHC (A) and western blot (B) of dog heart showing presence of GPR39. Arterioles show GPR39 in green. Nuclei are stained blue. Western blot depicts a characteristic 54 kDa band for GPR39 from 3 different dogs.

**Figure 2:**
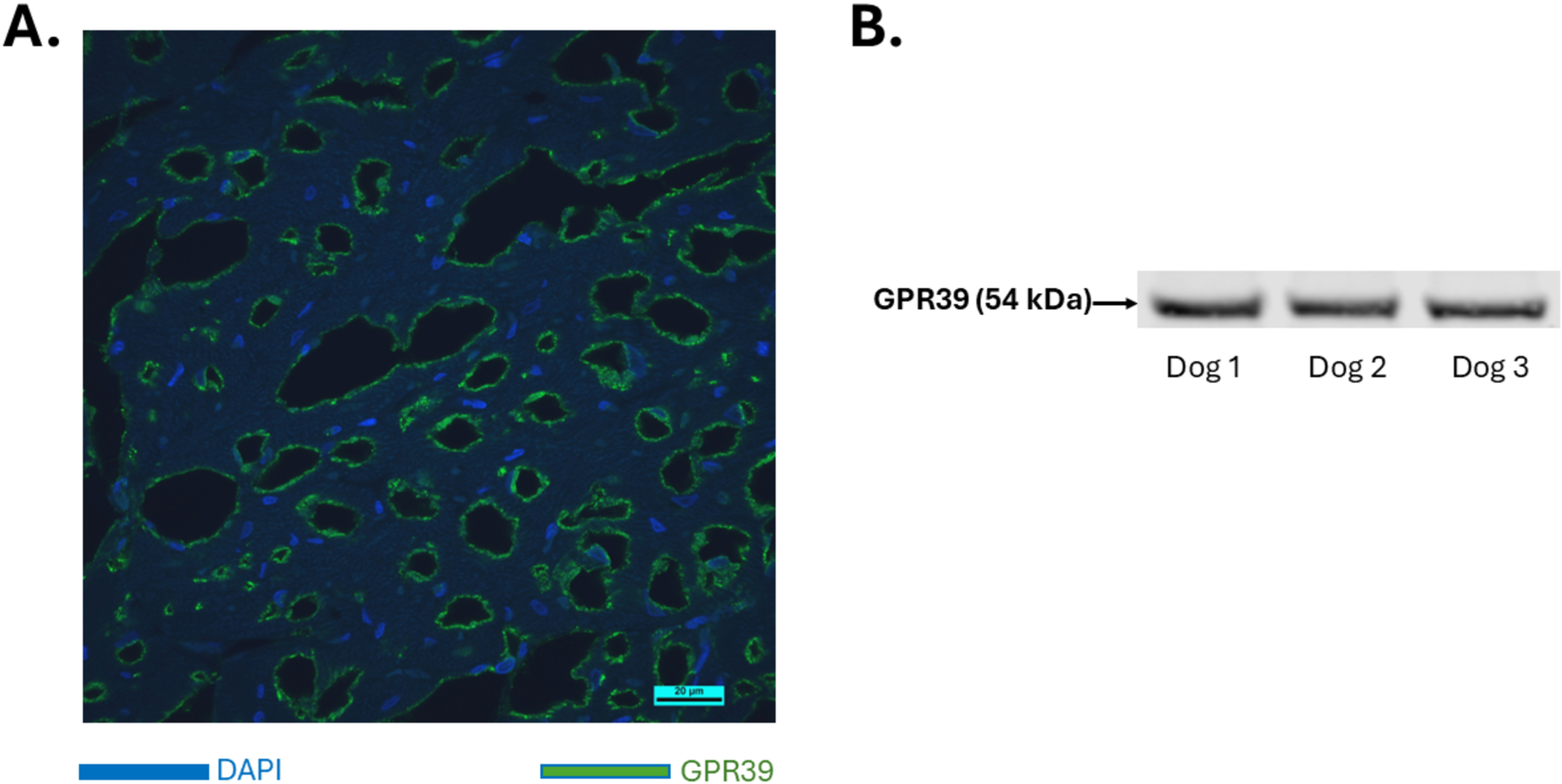
Examples of IHC (A) and western blot (B) of dog heart showing presence of GPR39. GPR39 (in green) is seen lining the arterioles (A). Nuclei are stained blue (DAPI). Western blot depicts a characteristic 54 kDa band for GPR39 from 3 different dogs. GPR39=G-protein coupled receptor, 39; DAPI=4′,6-diamidino-2-phenylindole

Table 1 lists the relation between CDP and all the eicosanoids in 6 Group 1 dogs during 17 stages, analyzed using linear mixed-effects models with a random intercept for dog and Bonferroni correction across the eicosanoid panel. After correction, 15-HETE (vasoconstrictor) and two dihydroxyeicosatrienoic acids (5,6-DHET and 8,9-DHET) that are vasodilators showed a significant relation with CDP: As CDP levels decreased so did 15-HETE (Figure 3A). When linear mixed model was repeated with only 15-HETE, 5,6-DHET and 8,9-DHET, the marginal r^2^ increased to 0.56 and only 15-HETE remained a significant predictor after adjusting for 5,6- DHETE and 8,9-DHET (p=0.0003). When adjusted for 15-HETE both 5,6-DHET and 8,9-DHET became non-significant (p=0.57 and p=0.44, respectively) with their coefficients becoming negative. 5,6-DHET and 8,9-DHET did not add significantly to the model fit over 15-HETE alone with the likelihood ratio test comparing the variables with the joint model (p=0.28). The levels of 14,15-EET (vasodilator) that acts through GPR39 did not change (Figure 3A). Since the ratio of 15-HETE (vasoconstrictor) and 14,15-EET (vasodilator) could influence coronary vascular tone through GPR39, we determined its relationship with CDP (Figure 3B) and, as expected based on our results, it remained significantly correlated with CDP (P=0.018).

**Figure 3:**
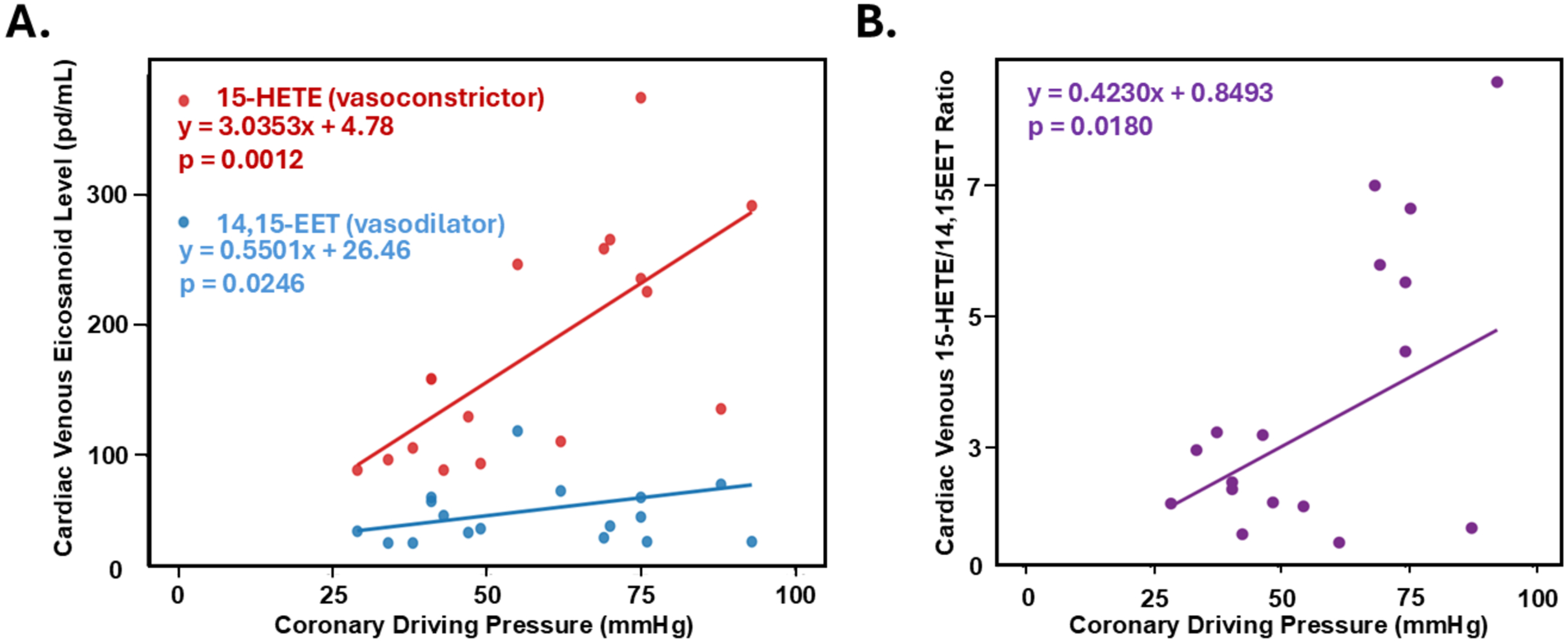
Relation between coronary driving pressure and anterior cardiac venous plasma levels of A)15-HETE and 14,15-EET from 7 dogs undergoing different degrees of coronary stenosis within the autoregulatory range, and B) 15-HETE/14,15-EET ratio. 15-HETE=15- hydroxyeicosatetraenoic acid; 14,15-EET=14,15-epoxyeicosatrienoic acid.

**Table 1.**
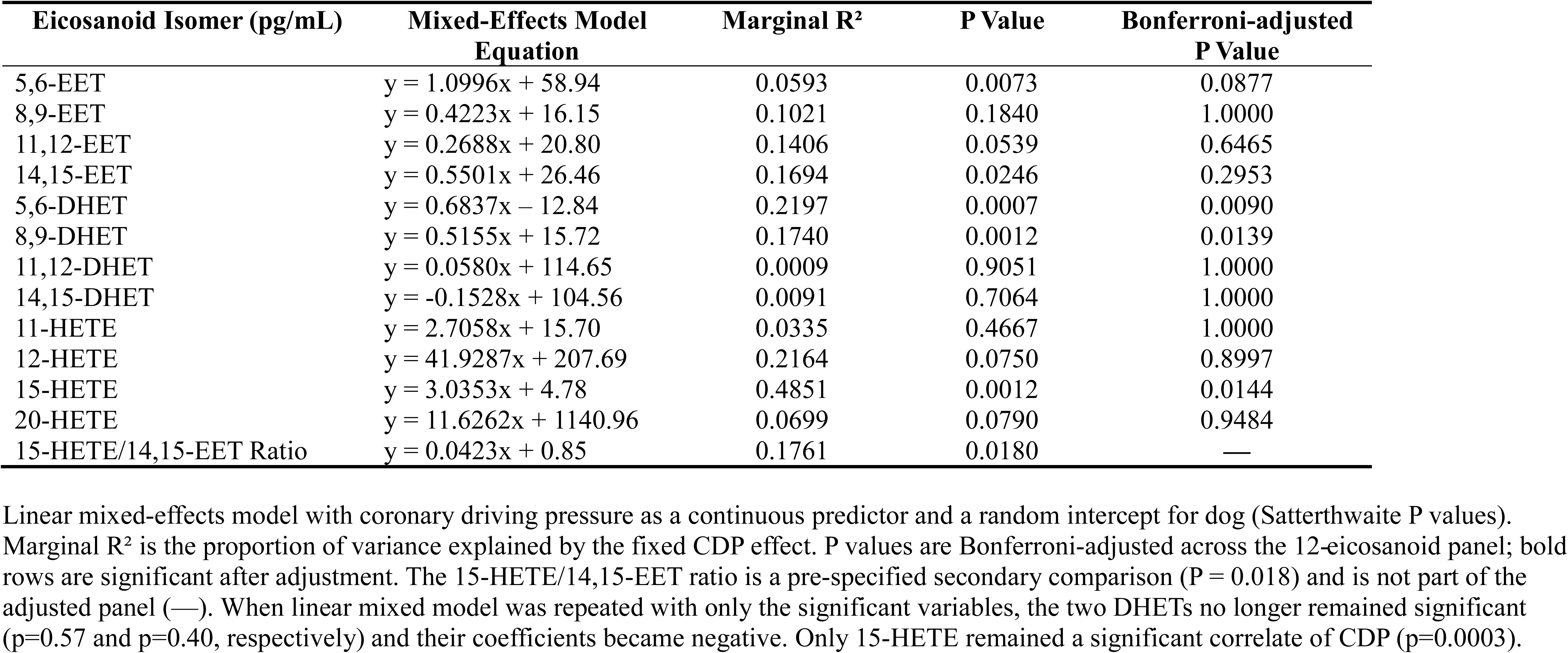
Relation between Coronary Driving Pressure and Coronary Venous Eicosanoid Levels (n = 6 dogs, 17 stages)

Since 15-HETE is the endogenous agonist for GPR39,^11^ we elected to test the effect of a specific GPR39 antagonist, VC108,^15^ on coronary autoregulation. This drug increases CBF in rodents,^16^ however, its hemodynamic effects in large animals are unknown. We measured its systemic, pulmonary, and coronary hemodynamics in addition to its effect on coronary autoregulation. Because various vasoactive metabolites have previously been implicated in the mechanism of coronary autoregulation,^9,10,17–22^ we performed comprehensive metabolomics of known coronary vasoactive compounds at different CDPs prior to VC108 administration.

We first created coronary stenoses (including where CDP declined below the autoregulatory range) of the left anterior descending coronary artery in 22 dogs and measured CDP and CBF during each stage (Figure 1). In 12 of these dogs, we sampled 2.5 mL of blood from the anterior cardiac vein for measurement of adenosine, endothelin-1, polyunsaturated fatty acids (including arachidonic acid), and prostaglandins. For prostaglandin and adenosine measurements blood was collected in stop solutions. We then removed the stenosis and allowed coronary hemodynamics to return to normal.

Table 2 lists the systemic and coronary hemodynamics at baseline and during different stenoses stages in 22 dogs (total number of stages = 77). Heart rate increased significantly after placement of severe stenosis. Systemic pressures did not change between stages. In contrast, as expected, CDP decreased with increasing degrees of stenoses and CBF remained constant for non-critical stenoses and declined when stenoses became severe (flow limiting). Constant CBF over a wide range of CDP with non-critical stenosis demonstrates that autoregulation was operative at baseline. MVR decreased progressively with increasing stenoses severity and increased with severe stenosis, although these changes were not statistically significant. We previously reported increased MVR with reduction in resting CBF despite exhaustion of autoregulation caused by a decrease in myocardial blood volume.^23^

**Table 2.**
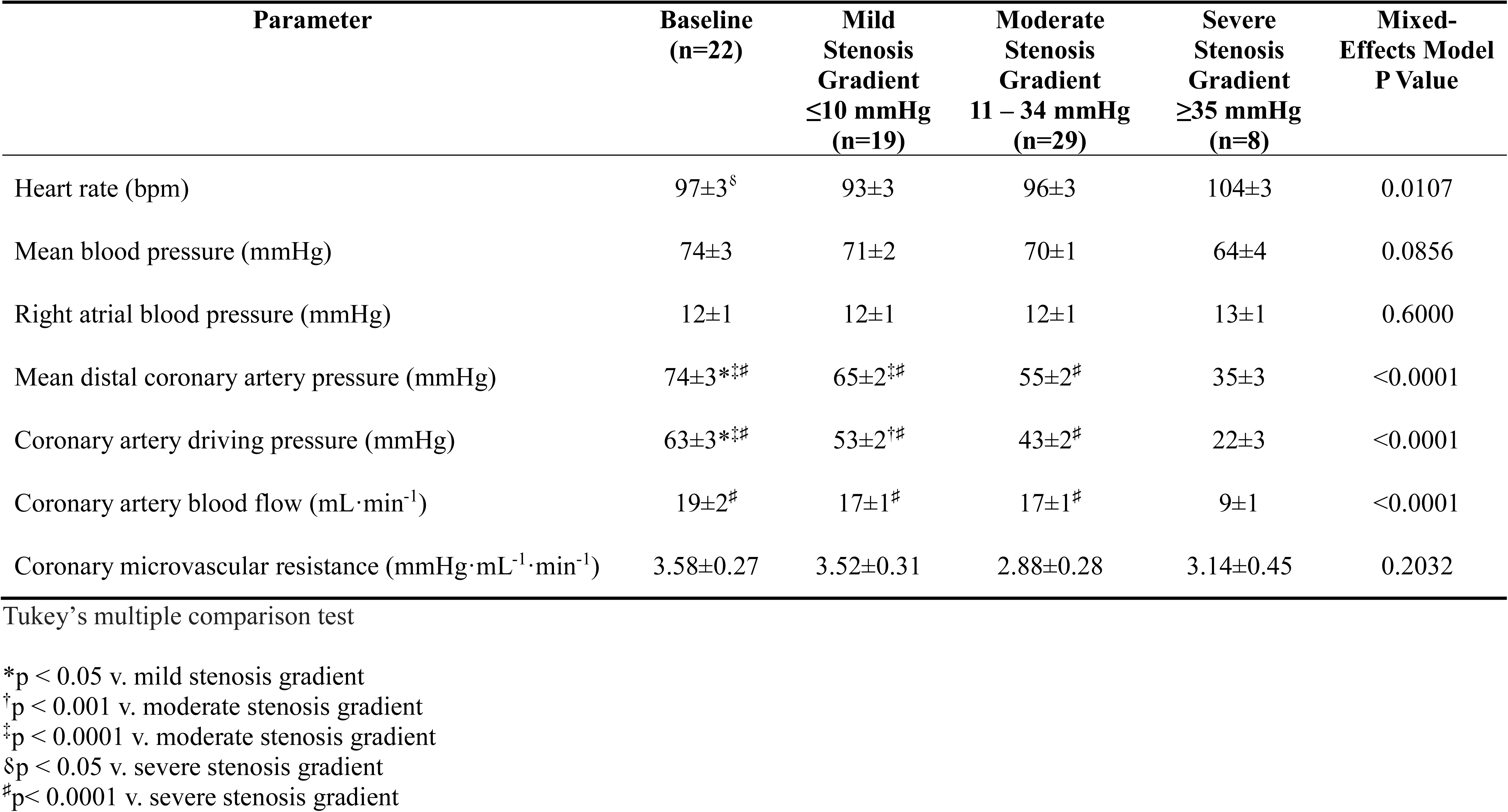
Systemic and Coronary Hemodynamics for 22 Protocol 3 Dogs at Baseline and Varying Degrees of Coronary Stenosis (n=77 Stages) to VC108 Administration (mean ± SEM)

| Parameter | Baseline<br>(n=22) | Mild<br>Stenosis<br>Gradient<br>≤10 mmHg<br>(n=19) | Moderate<br>Stenosis<br>Gradient<br>11 – 34 mmHg<br>(n=29) | Severe<br>Stenosis<br>Gradient<br>≥35 mmHg<br>(n=8) | Mixed-Effects Model<br>P Value |
| --- | --- | --- | --- | --- | --- |
| Heart rate (bpm) | 97±3<sup>§</sup> | 93±3 | 96±3 | 104±3 | 0.0107 |
| Mean blood pressure (mmHg) | 74±3 | 71±2 | 70±1 | 64±4 | 0.0856 |
| Right atrial blood pressure (mmHg) | 12±1 | 12±1 | 12±1 | 13±1 | 0.6000 |
| Mean distal coronary artery pressure (mmHg) | 74±3*<sup>‡</sup> | 65±2<sup>‡</sup> | 55±2<sup>#</sup> | 35±3 | <0.0001 |
| Coronary artery driving pressure (mmHg) | 63±3*<sup>‡</sup> | 53±2<sup>†</sup> | 43±2<sup>#</sup> | 22±3 | <0.0001 |
| Coronary artery blood flow (mL·min<sup>-1</sup>) | 19±2<sup>#</sup> | 17±1<sup>#</sup> | 17±1<sup>#</sup> | 9±1 | <0.0001 |
| Coronary microvascular resistance (mmHg·mL<sup>-1</sup>·min<sup>-1</sup>) | 3.58±0.27 | 3.52±0.31 | 2.88±0.28 | 3.14±0.45 | 0.2032 |

Tukey's multiple comparison test
\*p < 0.05 v. mild stenosis gradient
†p < 0.001 v. moderate stenosis gradient
‡p < 0.0001 v. moderate stenosis gradient
§p < 0.05 v. severe stenosis gradient

Figure 4A illustrates data from individual dogs in the presence of different degrees of coronary stenosis. There is wide variation in baseline CBF between animals depending on the size of the instrumented coronary artery. As a rule, CBF decreased only when CDP was low. Aggregate results from the 22 animals are shown in Figure 4B where autoregulation is clearly seen with mild and moderate degrees of stenosis, and CBF declined only when CDP was low (severe stenosis). The data fit well to a one phase decay function.

**Figure 4:**
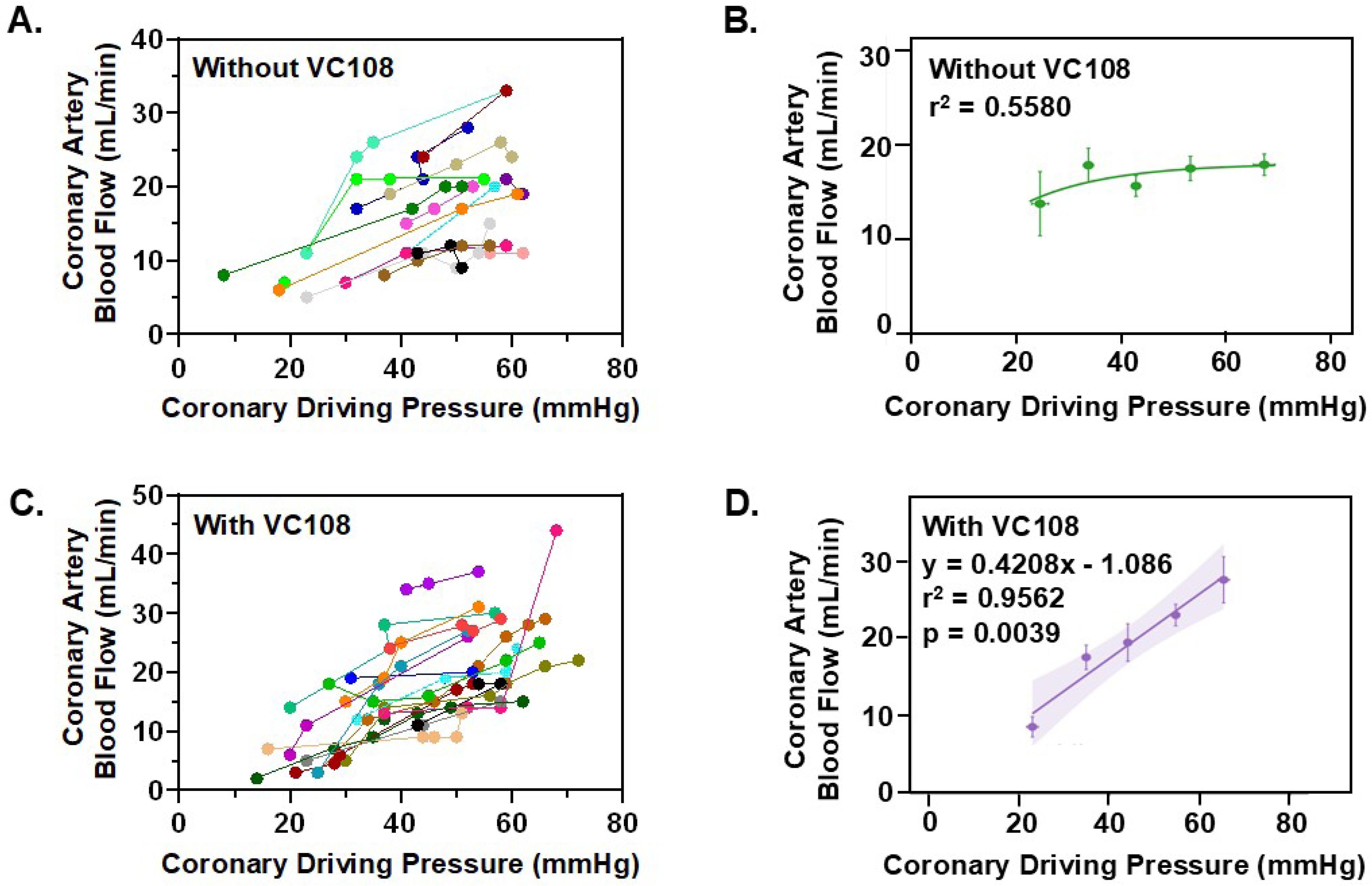
Coronary driving pressure versus coronary artery blood flow results from protocols 2 and 3. A) Data from 22 individual dogs (77 unique stages) in the absence of VC108 showing flat coronary artery blood flow over a wide range of coronary driving pressures, with decreasing flow at low coronary driving pressures. B) Aggregate results from the same animals as in panel A, showing classical coronary autoregulation over a wide range of coronary driving pressures with a decline in coronary artery flow at low coronary driving pressures. Data were fit using one phase decay least-squares analysis. C) Data from 14 of the 22 individual dogs (74 unique stages) from protocol 2 in the presence of VC108 (protocol 3) showing declining coronary artery blood flow with lowering of coronary driving pressure. D) Aggregate results from the same animals as in panel C, showing abolishment of coronary autoregulation with coronary artery blood flow becoming coronary driving pressure dependent. Data were fit using linear regression analysis.

Table 3 depicts the relation between CDP and different metabolites measured from the anterior cardiac vein during different stenosis stages. After correction for multiple comparisons, no cardiac venous metabolite correlated with CDP. Figure 5 illustrates the relation between CDP and adenosine and endothelin-1, while Figure 6 illustrates the relation between CDP and arachidonic acid and prostaglandins. These cardiac metabolites do not participate in coronary autoregulation.

**Figure 5:**
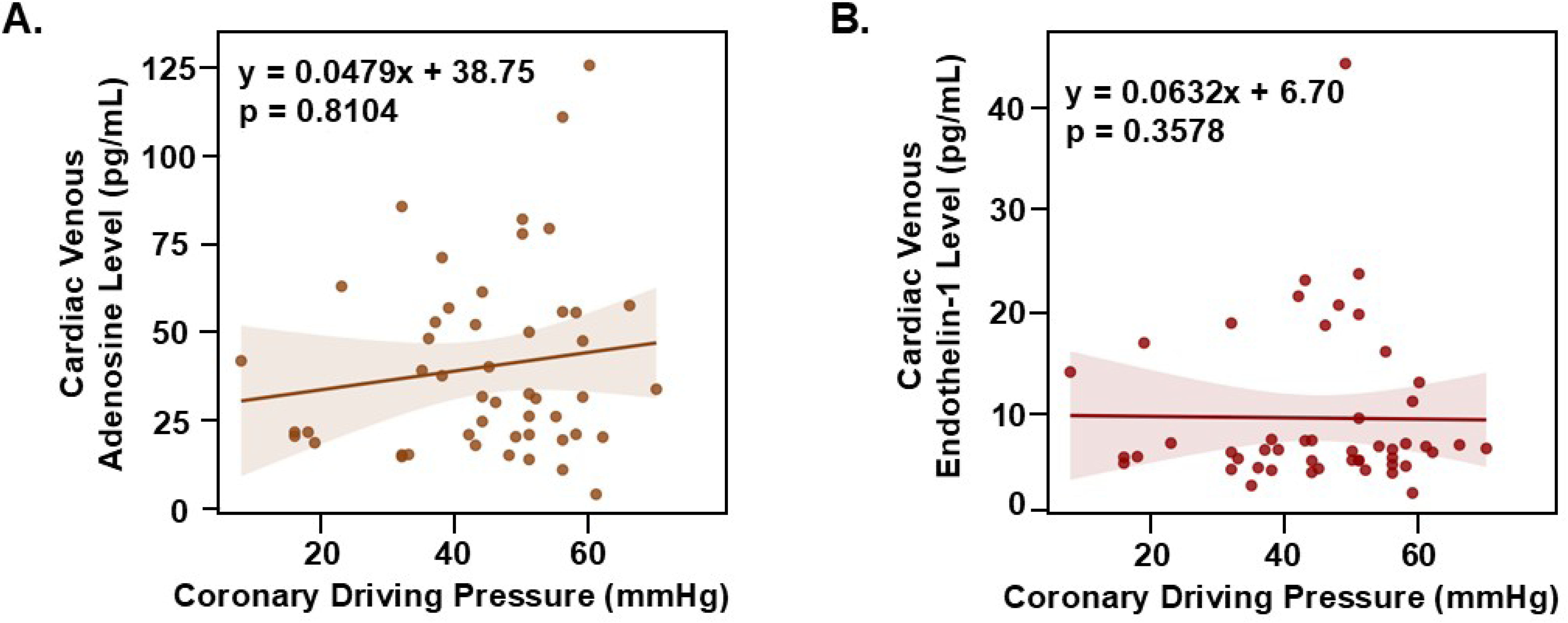
Relation between coronary driving pressure and anterior coronary vein plasma concentrations of A) adenosine and B) endothelin-1 from 12 dogs undergoing various degrees of coronary stenoses at rest.

**Figure 6:**
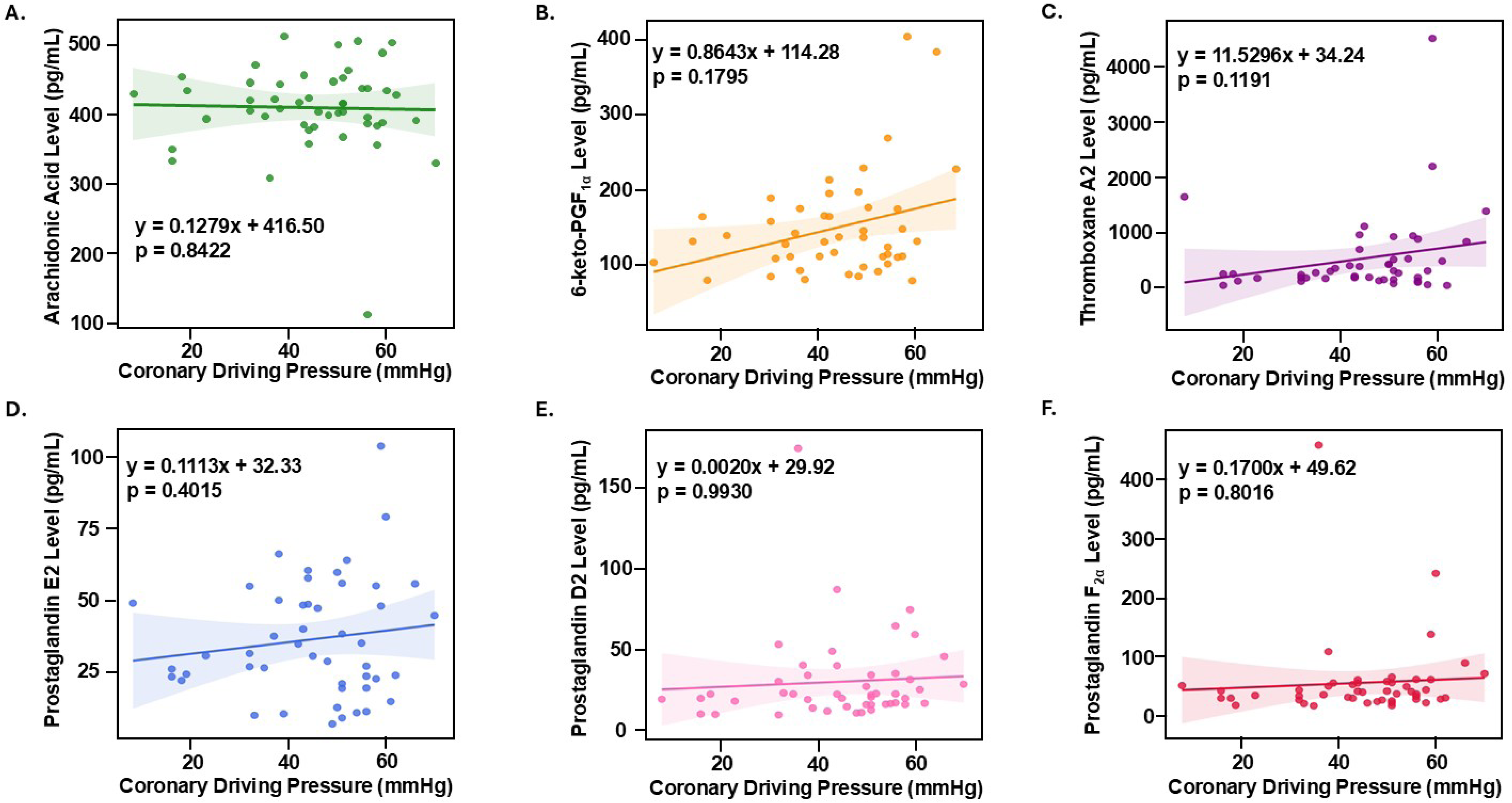
Relation between coronary driving pressure and anterior coronary vein plasma concentrations of A) Arachidonic acid; B) 6-keto-PGF_α_, (an inactive metabolite of prostaglandin I2; C) Thromboxane A2; D) Prostaglandin E2; E) Prostaglandin D2; and F) Prostaglandin F2_α_.

**Table 3.**
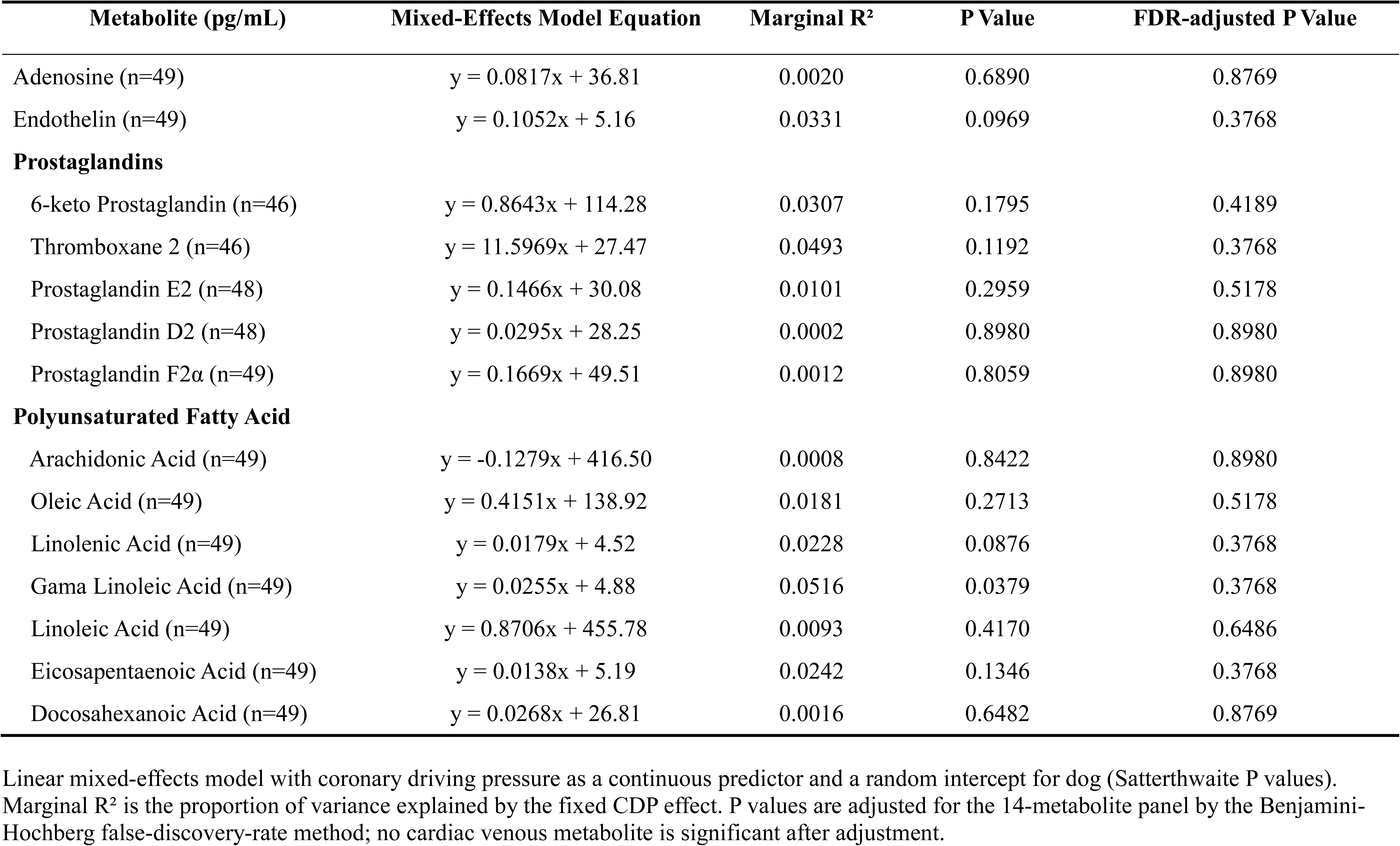
Relation between Coronary Driving Pressure and Cardiac Venous Metabolite Levels During Baseline and Various Stenoses Prior to 8 Administration in 12 Dogs.

After collecting these data, we initiated the administration of either VC108 (14 animals) or vehicle (8 animals). We added 3 more dogs to the vehicle group to obtain a comparable number to the VC108 group. Thirty minutes later we measured systemic, pulmonary, and coronary hemodynamics. Table 4 lists the baseline systemic and coronary hemodynamics before administration of VC108 (n=13) or vehicle (n=11). RA pressure was not measured in an additional dog receiving VC108 and data from it are not included in the table. At baseline, prior to administration of either vehicle or VC108, CDP was lower in dogs that were assigned to receive VC108 because of lower mean aortic and consequently distal coronary pressure.

**Table 4.** Baseline Systemic and Coronary Hemodynamics in Protocol 2 dogs Prior to Vehicle/Drug Administration (mean ± SEM)

| **Parameter** | **Vehicle**<br>**(n=9)** | **VC108**<br>**(n=10)** | **P value** |
| --- | --- | --- | --- |
| Heart rate (bpm) | 101±6 | 104±3 | 0.6092 |
| Mean aortic pressure (mmHg) | 80±5 | 72±2 | 0.1048 |
| Mean right atrial pressure (mmHg) | 11±2 | 12±1 | 0.5283 |
| *Left ventricular $\Delta P/\Delta t$ (mmHg·s$^{-1}$) | 1082±185 | 868±127 | 0.3533 |
| *Mean pulmonary artery pressure (mmHg) | 23±5 | 16±1 | 0.1461 |
| Distal coronary artery pressure (mmHg) | 82±4 | 72±2 | 0.0508 |
| Coronary artery driving pressure (mmHg) | 71±4 | 60±2 | 0.0108 |
| Coronary artery blood flow (mL·min$^{-1}$) | 21±2 | 22±2 | 0.9097 |
| Coronary microvascular resistance (mmHg·mL$^{-1}\cdot$min$^{-1}$) | 3.66±0.49 | 3.14±0.30 | 0.3747 |

\*Measured in 6 dogs

Figure 7 illustrates the change in hemodynamics after VC108/vehicle. There was no difference between VC108 and vehicle in terms of changes in systemic or pulmonary hemodynamics (panels A to E). In comparison, there was a small, albeit non-significant decrease in distal coronary artery pressure (CAP) caused by VC108 compared to vehicle (panel F) accompanied by a significant increase in CBF (panel G) resulting in a significant reduction in the calculated coronary microvascular resistance (MVR, panel H). These results indicate that at rest inhibition of GPR39 causes reduction in coronary MVR without any change in systemic and pulmonary vascular resistances. No change in cardiac work (product of heart rate, aortic pressure, left ventricular ΔP/Δt) was noted with either vehicle or drug.

**Figure 7:**
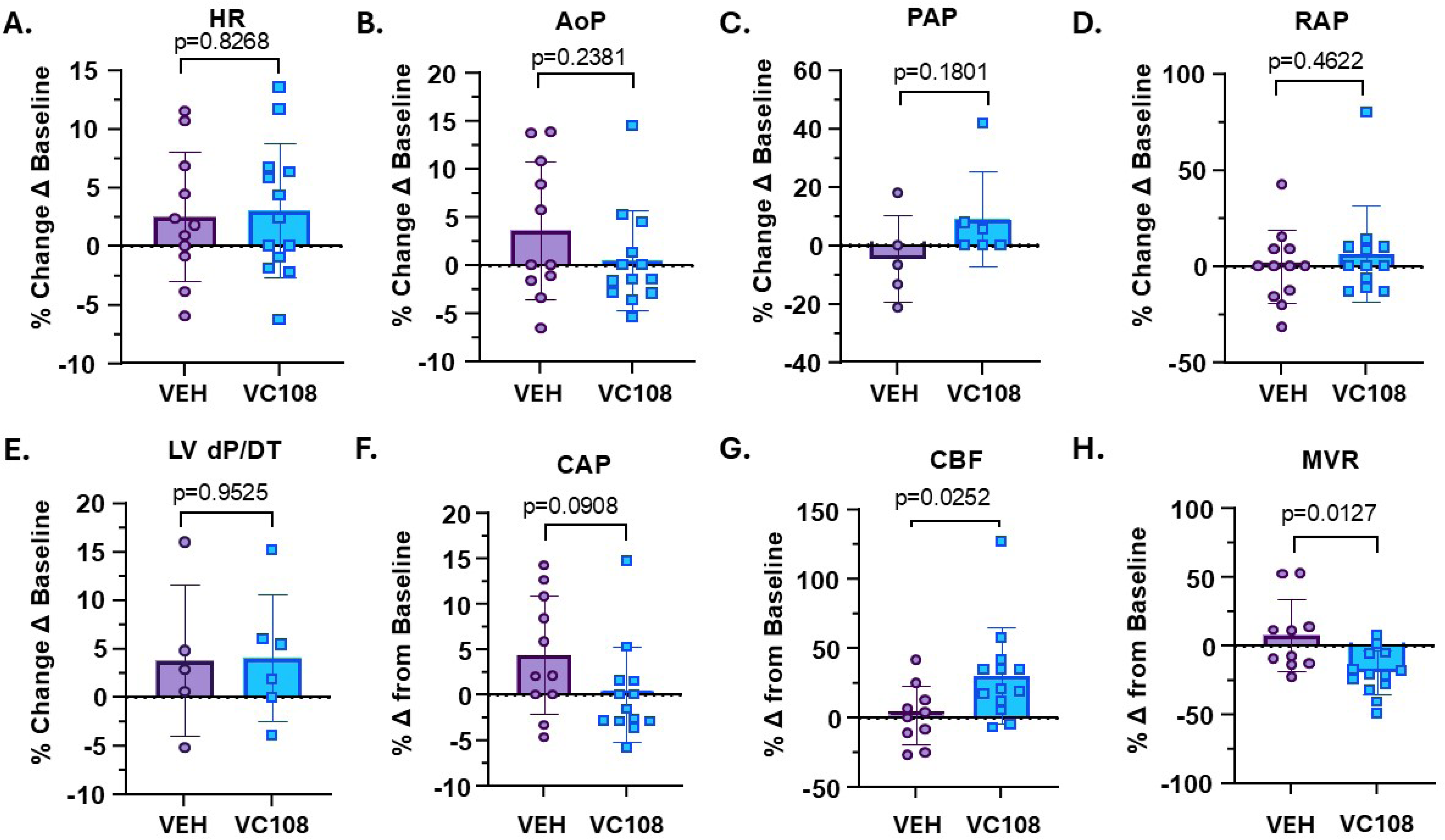
Percent change from baseline in systemic, pulmonary, and coronary hemodynamics with administration of Vehicle (VEH, n=11) or VC108 (n=13). A) HR=heart rate; B) AoP=aortic pressure; C) PAP=pulmonary artery pressure; D) RAP=right atrial pressure; E) LV dP/dt=rate of change of left ventricular pressure; E) CAP=coronary artery pressure; F) CBF=coronary blood flow; and H) MVR=coronary microvascular resistance. Comparisons between groups were made using unpaired *t-*test. **Central Illustration.** Mechanism of Coronary Autoregulation: Increased or decreased intraluminal pressure is sensed by ion channels or GPCRs that will either increase (high pressure) or decrease (low pressure) arachidonic acid release from plasma membrane. Arachidonic acid is then converted to leading to 15-HETE, the level of which determines coronary arteriolar tone. GPCR=G-protein coupled receptor. 15-HETE=15-hydroxyeicosatetraenoic acid; 15-LO=15- lipooxygenase; PLA_2_=phospholipase A_2_. Created in https://Biorender.com

We then recreated stenoses of varying severities in the 14 dogs receiving VC108 and measured CDP and CBF at each stage. Table 5 shows results from the different stenoses stages. Systemic pressures did not change between stages. Baseline CBF was higher (26±2 vs. 19±2 mL.min^-1^, p=0.05) and MVR was lower (2.42±0.20 vs. 3.58±0.27 mmHgꞏmL^-1^ꞏmin^-1^, p=0.03) during VC108 compared to prior to VC108 (Table 2). The increase in CBF compared to baseline was less than the maximal increase in CBF noted after 20 s coronary occlusion at the beginning of the experiment indicating that VC108 does not maximally dilate coronary arterioles. CBF declined with each level of non-critical stenosis, indicating abolishment of coronary autoregulation. Because of coronary vasodilation caused by VC108, MVR remained similar during all levels of non-critical coronary stenosis but increased when the stenosis was critical. We previously showed that the majority (>75%) of increase in MVR distal to critical stenosis during coronary vasodilation occurs from capillary derecruitment distal to stenosis.^23^

**Table 5.**
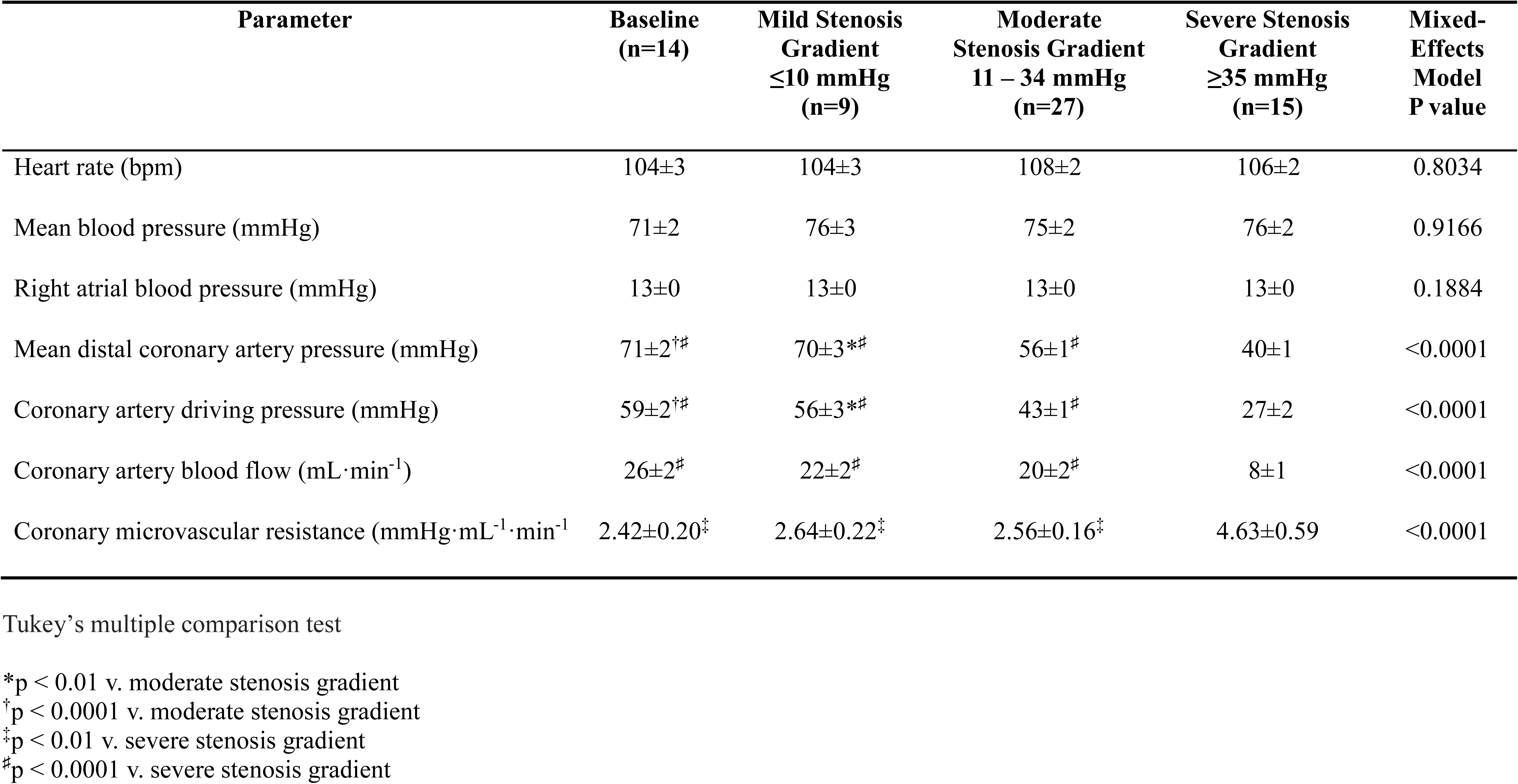
Systemic and Coronary Hemodynamics for 14 Protocol 3 Dogs at Baseline and with Varying Degrees of Coronary Stenoses (n=74 s) During VC108 Administration (mean ± SEM)

Figure 4C illustrates results from the 14 individual dogs where different degrees of stenoses were placed in the presence of intravenously administered VC108. As a rule, CBF decreased with decreasing CDP, although the slopes of the decline and baseline CBF varied between animals. Aggregate results of the 74 stages from 14 animals are shown in Figure 4D. There is a linear relation between CDP and CBF implying that VC108 abolishes autoregulation.

Table 6 illustrates a linear mixed-effects model comparison (condition-by-stage interaction with a random intercept for dog) of systemic and coronary hemodynamics during various degrees of stenosis in the absence (Table 2) or presence (Table 5) of VC108. Except for RA pressure, the condition-by-stage interactions were highly significant, indicating that hemodynamics changed differently across stenosis stages in the presence versus absence of VC108.

**Table 6.** Interactions between variables in. Tables 4 and 5 **(Protocols 2 and 3 Dogs)**

| **Parameter** | **F value** | **P value** |
| --- | --- | --- |
| Heart rate (bpm) | 21.22 | <0.0001 |
| Mean blood pressure (mmHg) | 52.22 | <0.0001 |
| Right atrial blood pressure (mmHg) | 0.98 | 0.4011 |
| Mean distal coronary artery pressure (mmHg) | 25.25 | <0.0001 |
| Coronary artery driving pressure (mmHg) | 23.65 | <0.0001 |
| Coronary artery blood flow (mL·min<sup>-1</sup>) | 30.55 | <0.0001 |
| Coronary microvascular resistance (mmHg·mL<sup>-1</sup>·min<sup>-1</sup>) | 86.41 | <0.0001 |

In 3 naïve dogs we administered VC108 and measured drug levels every 10 min for one hour (Table 7). The elevated level at 10 min is from the bolus injection (see methods), after which the drug level remains steady during VC108 infusion. The mean blood/plasma ratio from 3 animals was 0.98. Of note, the ratio of free plasma concentration/IC50 of VC108, which in the dog is 31.6 nM, is well over unity, implying therapeutic effect.

**Table 7.** Total (mean and range, nM) and Mean Free Blood and Plasma Concentration (nM) as well as Free Plasma Concentration/IC50 of VC108 After a 1.5 mg Bolus Injection Followed by 0.6 mg/Kg/h Infusion in 3 Protocol 1 Dogs.

| **Time After Administration**<br>**(min)** | **Total Blood**<br>**Concentration**<br>**(Range)** | **Free Blood**<br>**Concentration** | **Free Plasma**<br>**Concentration** | **Free Plasma**<br>**Concentration/IC50** |
| --- | --- | --- | --- | --- |
| **10** | 14752<br>(4054-31573) | 4661 | 4568 | 71 |
| **20** | 6824<br>(5234-8710) | 2156 | 2113 | 33 |
| **30** | 6382<br>(4664-8700) | 2017 | 1977 | 31 |
| **40** | 5876<br>(3900-8348) | 1857 | 1820 | 28 |
| **50** | 6094<br>(3618-10297) | 1926 | 1887 | 29 |
| **60** | 5902<br>(3767-8974) | 1653 | 1620 | 25 |

Blood plasma ratio of VC108=0.98
IC50 of VC108=31.6 nM

In addition, the 18 Group 3 beagle dogs (9 female and 9 male) were administered daily intravenous bolus injections of VC108 at 3 different doses (2, 6, and 20 mgꞏkg^-1^) and pharmacokinetics were determined at days 1 and 14. There was no difference in drug exposure between female and male dogs at all 3 doses both on day 1 and day 14 (Supplemental Results). Of the 131 receptors, channels, and enzymes studied for off-target binding, VC108 did not bind to any of them (Supplemental Results).

## Discussion

Using a comprehensive metabolomic analysis of vasoactive molecules that could have a putative role in coronary autoregulation, we show for the first time that coronary venous levels of 15-HETE, the endogenous agonist of GPR39, correlates with changes in CDP, whereas other metabolites do not. Since 15-HETE acts through GPR39, our results indicate that GPR39 is the molecular switch that controls coronary vasomotion at rest. This is supported by the finding that after administration of VC108, a specific GPR39 inhibitor, coronary MVR decreased. In the absence of VC108, CBF remained constant over a wide range of CDPs induced by varying degrees of coronary stenosis indicating intact autoregulation. VC108 abolished coronary autoregulation so that CBF became CDP dependent. Therefore, GPR39 and its endogenous agonist 15-HETE together orchestrate coronary autoregulation.

In our previous study, where we showed that GPR39 was the receptor for the endogenous ligands 15-HETE and 14,15-EET in the mouse, we stripped the heart of all large vessels and demonstrated the presence of GPR39 on VSMCs in the coronary microvasculature using immunohistochemistry: GPR39 was present in coronary resistance arterioles^11^, similar to results from our current study in dogs. In addition, we showed the presence of GPR39 in mouse heart using western blot^11^, similar to results from our current study in dogs. Using a mouse Langendorf constant flow preparation we showed that 15-HETE increased CDP above the normal range, an effect that was abolished in GPR39 knockout mice. In these experiments, 14,15-EET alone had no effect on CDP. Our current results extend these observations.

GPR39 exhibits constitutive activity through Gαq and Gα12/13, which may explain its role in maintaining resting coronary vascular tone.^24,25^ Zn^++^ serves as an allosteric modulator of GPR39.^24,25^ In the presence of Zn^++^, GPR39 activation by 15-HETE leads to extracellular signal-regulated kinase (ERK) phosphorylation and intracellular Ca^++^ mobilization,^11^ suggesting activation of Gαq. Increased intracellular Ca^++^ results in VSMC contraction. The higher the level of 15-HETE, the greater the stimulation of GPR39 and the greater the intracellular Ca^++^ and consequent vasoconstriction.^11^ As 15-HETE levels decrease the reverse occurs, leading to less intracellular Ca^++^ and less vasoconstriction (or more vasorelaxation)

### Role of Arachidonic acid Metabolites in Coronary Autoregulation

Prostaglandins, another set of arachidonic acid derivatives, were examined previously where administration of two inhibitors of prostaglandin synthesis (5,9,11,14-eicosatetraynoic acid and indomethacin) abolished coronary autoregulation in the perfused isolated rabbit heart,^26^ although studies in larger animals did not show any effects of indomethacin administration on CBF.^27,28^ The same investigators who reported the effect of prostaglandin synthesis inhibition in the isolated rabbit heart subsequently reported an ‘active’ substance, an unidentified albumin bound lipid molecule, that counteracted the effects of indomethacin.^29^ Although it cannot be definitively proven, our results suggest that this ‘active’ substance could have been 15-HETE, a metabolite of arachidonic acid, which the investigators did not measure. When human endothelial cells are stimulated by histamine, they enhance synthesis of both prostaglandin and 15-HETE.^30^ Both are inhibited by indomethacin, which might explain why autoregulation was abolished with the drug in the perfused isolated rabbit heart.^31^ Our results indicate that prostaglandins, specifically TBXA2, PDE2, PGE2, and PGF2_α_, do not participate in autoregulation.

Of the eicosanoids, another group of arachidonic acid metabolites, only 15-HETE and two dihydroxyeicosatrienoic acids (5,6-DHET and 8,9-DHET) showed a significant correlation with CDP in our study. 5,6-EET is unstable and converts rapidly to 5,6-DHET. The δ-lactone of 5,6- DHET is more stable than 5,6-DHET and has been shown to be a potent coronary vasodilator in the dog. So has 8,9-DHET.^22^ Specific receptors for these metabolites have not been defined. However, when adjusted for 15-HETE, 5,6-DHET and 8,9-DHET no longer correlated with CDP and their coefficients became negative, which makes physiological sense since CDP reduction should be associated with an increase in vasodilator levels. The levels of 14,15-EET (vasodilator) did not change (Figure 3A). Coronary venous plasma levels of arachidonic acid itself remained unchanged at different CDPs, indicating that changes in coronary pressure influenced arachidonic acid to 15-HETE conversion by 12/15 LOX probably by release of more substrate from the lipid bilayer by phospholipase A_2_ (PLA_2_).

### Role of Other Pathways-Comparison with Previous Studies

Nitric oxide (NO) has been implicated in coronary autoregulation.^17^There are several pathways through which GPR39, 15-HETE, and NO interact with each other to explain these findings. First, increased intracellular Ca^++^ stimulates endothelial NO synthetase (eNOS) resulting in increased NO production.^32^ NO, in turn, stimulates sarcoplasmic endoplasmic reticulum Ca^++^ATPase, thereby increasing Ca^++^ reuptake into sarcoplasmic endoplasmic reticulum and decreasing vascular tone.^33^ GPCRs can undergo post-translation modification induced by S-nitrosylation whereby NO could modify the response of GPR39 to 15-HETE in vivo.^34^ Thus, inhibition of eNOS by NG-nitro-L-arginine could interfere with autoregulation as previously reported^17^ by inhibiting signaling both at the level of GPR39 and downstream to it. However, we did not test this possibility in our current study.

Inhibition of α-adrenergic systemic has been shown to alter coronary autoregulation.^18^ α- adrenergic stimulation causes increased production of arachidonic acid through activation of mitogen-activated protein kinases that are downstream to ERK, enhancing arachidonic acid production from the nuclear envelope.^35^ It also increases production of arachidonic acid and its metabolites by increasing intracellular Ca^++^, which stimulates cytosolic PLA_2_ production that is responsible for release of arachidonic acid from plasma membrane.^36^ Thus, α-adrenergic inhibition by drugs such as prazocin reported previously^18^ could affect coronary autoregulation through inhibition of arachidonic acid and 15-HETE production, mechanisms proximal to GPR39.

Vasodilation requires VSMC hyperpolarization that is mediated through increased levels of intracellular K^+^ resulting from K^+^ channel opening.^19^ 15-HETE directly inhibits K^+^ channel activity, thus attenuating vasodilation.^37^ Lower levels of 15-HETE seen with coronary arteriolar vasodilation at lower CDPs are therefore likely associated with more active K^+^ channels that are open and allow intracellular K^+^ flux under appropriate circumstances. Open K^+^ channels-induced hyperpolarization also decreases Ca^++^ sensitivity.^38^ Other K^+^ channel blockers have also been associated with disruption of coronary autoregulation.^10,20,39^ H_2_O_2_ opens Ca^++^ gated and voltage-dependent K^+^ channels allowing VSMC hyperpolarization.^39^ Therefore, K^+^ channel blockers like tetraethyl ammonium and 4-aminopyridine block the effect of H_2_O_2_, thus interfering with vasodilation.^20^ A similar explanation can be provided for attenuation of coronary autoregulation by glibenclamide, an ATP-sensitive K^+^ channel blocker.^20^ K^+^ channel activity is downstream to GPR39 activation.

Finally, adenosine was previously shown not to participate in coronary autoregulation^9,10^ and we confirmed the finding. Additionally, we show that endothelin-1, a potent coronary vasoconstrictor^40^ also does not participate in coronary autoregulation.

### Proposed Mechanism of Coronary Autoregulation

Based on our results we propose the following scenario for coronary autoregulation depicted in the central illustration. Changes in CDP are mechanosensed by ion channels^41,42^ or G- protein coupled receptors^43–45^ present on VSMCs, which then either activate PLA_2_ to release the arachidonic acid from plasma membrane to produce its metabolites or directly modulate 12/15 LOX activity resulting in 15-HETE production. 15-HETE then activates GPR39, which in turn causes intracellular Ca^++^ release resulting in VSMC contraction. Higher CDP causes higher 15- HETE production and more vasoconstriction while lower CDP causes less 15-HETE production and less vasoconstriction (or relative vasodilation). Counterintuitively, no vasodilators actively participate in this process. Coronary vascular tone needs to be rapidly adjusted to myocardial oxygen needs or CDP changes, implying that the molecular switch for coronary autoregulation needs to act more like a thermostat than a lock and key, which is readily accomplished by a single biomolecule acting on a single receptor. Having the same receptor detect changes in ambient arteriolar pressure would make the autoregulatory circuit even more efficient.

### Limitations of our Study

Sex differences are well known to affect coronary vascular reactivity in dogs.^46,47^ Uncertainty of the time during the ovulation cycle or possible pregnancy, prevented us from using female dogs. Thus, our results cannot be extrapolated to females in terms of vascular reactivity at baseline and during VC108 administration. In terms of exposure of animals to VC108 we found that both sexes of Beagle dogs had equal exposure to the drug.

Anesthesia can also affect coronary vascular reactivity. Although there are no direct interactions reported between the combination of hydromorphone, ketamine, and midazolam with arachidonic acid metabolism and GPR39 activation, ketamine indirectly modulates the arachidonic acid pathway by inhibiting the N-methyl-D-aspartate receptor-mediated calcium influx^48^, resulting in reduced PLA_2_ activity and arachidonic acid release. However, this effect likely did not confound our results as arachidonic acid levels were not affected throughout the range of CDP that was studied.

We did not study the upper range of autoregulation (100-120 mm CDP). Nonetheless, based on our previous results using a constant flow Langendorff mouse model,^11^ we predict that increasing CDP above the normal range would increase 15-HETE levels and cause further vasoconstriction to maintain constant CBF. In the dog, intracoronary injection of 15-HETE causes CBF decline from increased MVR that is mitigated by pre-treatment with VC108.^49^ We did not see elevation of adenosine or endothelin-1 levels at low CDP probably because ischemia was not produced due to adequate collateral flow to the myocardial region supplied by the left anterior descending artery. Collateral blood flow is high in the dog.^50^

While 15-HETE is the main metabolite that showed a correlation with CDP, the r^2^ value is <50 with significant scatter, which may be related to collateral blood flow, variability in coronary anatomy between dogs, and measurement variability. In addition, since CBF is highly regulated, other redundant and overlapping factors (neural, metabolic, etc.) may also contribute to coronary autoregulation.

Our results relating to GPR39 signaling are from either human prostate cancer cells that overexpress the protein^15^, transfected human embryonic kidney cells^11^, or VSMCs from mice.^11^ Whether canine GPR39 behavior is akin to that in these model systems is not known. Whereas >80-90% GPR39 homology between vertebrate species has been reported^51,52^, to our knowledge it has not been specifically studied in dogs. The G-proteins that are activated after GPR39 stimulation are similar for most species except fish.^51^

We did not measure cardiac metabolite levels during VC108 administration because in our early experiments we noted hemolysis after centrifugation of the coronary venous blood samples. It was then determined that hemolysis was caused by DMSO.^53^ Since then, additional pre-clinical studies have been performed using polyethylene glycol and cyclodextrin as excipients, without any in-vitro hemolysis. Development of the drug for clinical use will utilize as yet undetermined solvents.

GPR39 has been reported to have significant constitutive activity via its Gα_q_ and Gα_12/13_ subunits^54,55^, which may contribute to baseline coronary tone. We have previous evidence that VC108 blocks Gαq^15^ but have not yet studied its effects on Gα12/13. Not knowing the 15-HETE levels during different degrees of coronary stenoses after VC108 administration precluded studying the feedback loop in terms of GPR39 activation or desensitization. Whereas most agonists cause downregulation of G-proteins via β-arrestin^56^, GPR39 is unique in that it requires rho-kinase for desensitization.^57^ Antagonists like VC108 do not cause desensitization and generally cause G-protein coupled receptor upregulation.

### Conclusion

In this study we show for the first time that 15-HETE participates in the resting coronary vascular tone in dogs. We also demonstrate that 15-HETE acting through GPR39 is responsible for coronary autoregulation when CDP is reduced. Factors that have previously been described to effect coronary autoregulation either modify GPR39 directly or act through signaling pathways upstream or downstream of GPR39. Hence, GPR39 serves as the molecular switch responsible for coronary autoregulation.

## Supporting information

https://www.dropbox.com/scl/fi/nciovwsxq87ht1h5sp6le/Supplemental-material.pdf?rlkey=qyhl7w8uc639otqowdghgijhm&dl=0

## Non-standard Abbreviations and Acronyms

14,15-EET: 14,15-epoxyeicosatrienoic acid
15-HETE: 15-hydroxyeicosatetraenoic acid
CBF: coronary blood flow
CDP: coronary driving pressure
COX-1: cyclooxygenase-1
DHET: Dihydroxyeicosatrienoic acid
DMSO: dimethyl sulfoxide
EDTA: Ethylenediaminetetraacetic acid
eNOS: endothelial NO synthetase
ERK: extracellular signal-regulated kinase
Gαq: G-protein αq
GPR39: G-protein coupled receptor
H_2_O_2_: hydrogen peroxide
LOX: lipoxygenase
MVR: microvascular resistance
NO: nitric oxide
PGD2: prostaglandin D2
PGE2: prostaglandin E2
PGF2α: prostaglandin F2α
PLA2: phospholipase A2
RA: right atrial
TBXA2: thromboxane A2
VSMC: vascular smooth muscle cell

## Author Contributions

Drs. Le and Kaul developed the concept and design for the studies. They also wrote the manuscript. Drs. Kajimoto and Zhao participated in the studies and performed the surgery and delivered anesthesia and drugs. Dr. Methner performed ELISA for endothelin-1, western blot for GPR39, and prepared VC108 bolus and infusions. Dr. Cao performed immunohistochemistry. Dr. Le created the tables and figures. Dr. Minner performed the statistical analysis. Drs. Cianciulli, Semeraro, Trist and Micheli contributed to the development of VC108. Drs. Franchi, Marcheselli, and Parazzoli performed the pharmacokinetic studies in beagle dogs.

## Acknowledgements

Metabolomic analysis was performed by the Bioanalytical Shared Resource/ Pharmacokinetics Core Facility, which is supported in part by the University Shared Resource Program at Oregon Health and Sciences University.

The research reported in this publication used computational infrastructure supported by the Office of Research Infrastructure Programs, Office of the Director, of the National Institutes of Health under Award Number S10OD034224.

We acknowledge Biorender.com where the central illustration was crafted.

## Funding Sources

The Garthe and Grace L. Brown Fund and the John E. and Robin Jaqua Fund at the Oregon Community Foundation, and the Ernest C. Swigert endowment at the Oregon Health & Science University.

## Disclosures

Oregon Health & Science University and Vasocardea, Inc. have a joint global patent (WO2021222858 and corresponding national patents) that encompasses drugs that inhibit GPR39. Vasocardea, Inc. has an additional patent application filed globally (see WO2023076219 and corresponding national filings) that encompasses VC108. Vasocardea, Inc. has exclusive rights to develop and commercialize drugs under these patents. Evotec has assigned all patent rights to Oregon Health & Science University and Vasocardea, Inc.

Dr. Kaul is the founder and President of Vasocardea, Inc., a Delaware incorporated company located in Portland, Oregon.

## Disclaimer

The content is solely the responsibility of the authors and does not necessarily represent the official views of the Department of Veterans Affairs or the US Government.

## Supplemental Material

Supplemental Methods; Supplemental Results, Supplemental Table 1.

---

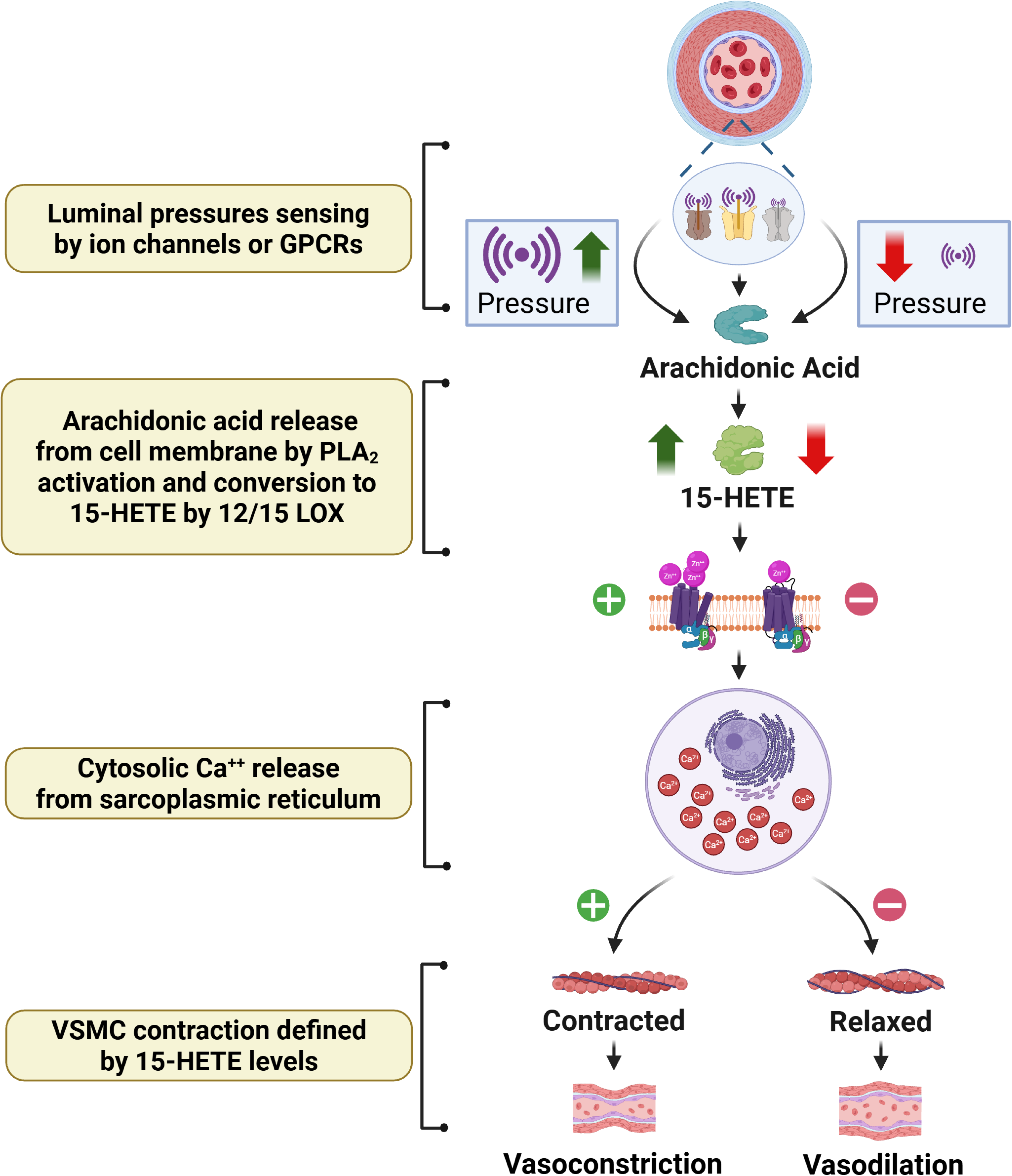

## Notes

### Competing Interest Statement

Oregon Health & Science University and Vasocardea, Inc. have a joint patent application filed globally (see WO2021222858 and corresponding national filings) that encompasses drugs that inhibit GPR39. Vasocardea, Inc. has an additional patent application filed globally (see WO2023076219 and corresponding national filings) that encompasses VC108. Vasocardea, Inc. has exclusive rights to develop and commercialize drugs under all of these patent applications. Evotec has assigned all patent rights to Oregon Health & Science University and Vasocardea, Inc. Dr. Kaul is the founder and President of Vasocardea, Inc., a Delaware incorporated company located in Portland, Oregon. The information in this manuscript does not represent the views of the Veterans Administration or the US government. ?

### Summary of Updates

Paper has been revised using a different statistical approach requested by the reviewers. Additional inofrmation of GPR39 has been added

