## Supplementary material for "15-Hydroxyeicosatetraenoic Acid and GPR39 Together Orchestrate Coronary Autoregulation: A Comprehensive Metabolomic Analysis": https://www.dropbox.com/scl/fi/nciovwsxq87ht1h5sp6le/Supplemental-material.pdf?rlkey=qyhl7w8uc639otqowdghgijhm&dl=0

**Supplemental Methods**

**Pharmacokinetics of VC108 in the Beagle Dog**

*Aim*: The objective of this study was to assess the pharmacokinetics of VC108 in the Beagle dog following daily intravenous administration for 14 days

*Methods*: VC108 was given to dogs (3/sex/group) at 2, 6 and 20 mg/kg/day once daily for 14 days by intravenous (bolus) administration. Dose volume administered was 2.5 mL/kg/day. Pharmacokinetic parameters were evaluated on Days 1 and 14.

**Receptors, Enzymes, Channels, and Uptake Binding of VC108**

*Aim*: To test VC108 binding on a panel of assays including receptors, enzymes, channels. and transporters.

*Methods*: VC108 was tested at 1.0E-05 M. Compound binding was calculated as a % inhibition of the binding of a radioactively labeled ligand specific for each target. Compound enzyme inhibition effect was calculated as a % inhibition of control enzyme activity.

**Supplemental Results**

**Pharmacokinetics of VC108 in the Beagle Dog**

On day 14 after repeated administration, mean maximal observed plasma concentration of VC108 (C_max_) increased with no major deviation from proportionality across the entire dose range in both male or female dogs, while the area under the plasma concentration time curve from the time of dosing to the last measurable concentration (AUC_last_) increased supra-proportionally from 2 to 6 mg/kg/day and in approximate proportion from 6 to 20 mg/kg/day in males and across all dose range in females. The supra-proportionality observed in AUC_last_ from 2 to 6 mg/kg/day in males is a consequence of a notable reduction in AUC_last_ at 2 mg/kg/day, due to increased clearance and faster elimination (decrease in terminal half-life) of VC108 after repeated administration compared to Day 1. No notable (>2-fold) differences in C_max_ were observed between single and repeated administration across the dose range in both sexes and in AUC_last_ in females at 2 mg/kg/day and in both sexes at 6 and 20 mg/kg/day. The exception was one female at 2 and 20 mg/kg/day, respectively) in which a notable decrease in AUC_last_, was associated with a decrease in terminal half-life and increase in clearance after repeated administrations. A nonsignificant decrease in AUC_last_ was observed after repeated administration also in two males and in one female given 20 mg/kg/day. No notable gender differences in systemic exposure (as mean C_max_ and AUC_last_) were observed across the dose range after both single and repeated administration (supplemental Table 1).

**Receptors, Enzymes, Channels, and Uptake Binding of VC108**

Of 131 receptors, channels, transporters and enzymes, including those involved in the cardiovascular system, only V1a(h) gave a 72% response in binding. The assay on this target was repeated to test functional activity of VC108 both in agonist and antagonist mode. VC108 gave no functional response. Therefore, VC108 is also functionally inactive at the V1a receptor.

We studied the effects of VC108 on C3 stimulation of GPR39 in human prostate cancer cells that overexpress GPR39. C3 is a potent synthetic agonist of GPR39. We found that we could inhibit C3 induced GPR39 stimulation with VC108 in a dose-dependent manner. Interestingly, mouse VSMCs produced very similar results.

**Supplemental Table 1 (Results of Pharmacokinetic Studies)**

| **Dose (mg/kg/day)** | **Male (3 animals at each dose)** | | | | **Female (3 animals at each dose)** | | | |
| --- | --- | --- | --- | --- | --- | --- | --- | --- |
|  | **Mean C_max_**  **(ng/mL)** | | **Mean AUC_last_**  **(ng∙h/mL)** | | **Mean C_max_**  **(ng/mL)** | | **Mean AUC_last_**  **(ng∙h/mL)** | |
|  | **Day 1** | **Day 14** | **Day 1** | **Day 14** | **Day 1** | **Day 14** | **Day 1** | **Day 14** |
| 2 (3 M, 3 F) | 2320 | 1740 | 2200 | 890 | 2060 | 1700 | 2120 | 1490 |
| 6 (3 M, 3F) | 7920 | 7370 | 8230 | 9360 | 6930 | 6640 | 6260 | 6260 |
| 20 (3 M, 3 F) | 27900 | 24200 | 45400 | 28200 | 22900 | 15800 | 29500 | 20600 |
